# Histamine signaling *via* the metabotropic receptor *Trapped in endoderm 1* regulates courtship initiation in *Drosophila melanogaster*

**DOI:** 10.1101/150680

**Authors:** Sadaf A. Zaki, Jaspal Sandhu, Rachael L. French

**Affiliations:** Department of Biological Sciences, San José State University, 1 Washington Square, San José, CA 95192-0100

## Abstract

Understanding the role of genes in directing behavior is one of the primary goals of neuroscience. Mating behavior in *Drosophila* is controlled by male-specific splicing of the master regulatory gene *fruitless* (*fru*), and the male-specific splice form, *fru^M^*, is both necessary and sufficient for all aspects of the courtship ritual. We have previously described the role of *Trapped in endoderm 1* (*Tre1*) in courtship behavior. *Tre1* encodes an orphan G-protein-coupled receptor that is essential for normal courtship behavior in male flies. We previously found that feminizing *Tre1*-expressing cells in males via expression of the female-specific splicing factor Transformer (Tra^F^) resulted in rapid courtship initiation. Here we confirm that Tre1 is required in neurons for normal courtship behavior, and present genetic evidence that Tre1 acts through the downregulation of the E-cadherin Shotgun, and that the neurotransmitter histamine is the likely Tre1 ligand. Our findings are the first evidence for metabotropic histamine receptors in *Drosophila*, and the first to demonstrate a role for histamine in courtship.

## Introduction

Genetic regulation of behavior is a vital topic in neuroscience—understanding gene-controlled behaviors will help to elucidate the intricacies of the central nervous system (CNS). The common fruit fly *Drosophila melanogaster* is an ideal model organism for the study of the genetic control of behavior. *Drosophila* has been used as a genetic model organism for over 100 years and provides a vast array of genetic manipulation techniques, complex and instinctive innate behaviors, and simple methods for observation of neuronal activity.

Male *Drosophila* carry out a complex courtship ritual involving a specific sequence of steps which depend on the processing of sensory cues by the CNS. First, the male fly orients himself toward the female fly and follows her. He then taps her with his forelegs, which contain gustatory receptors that he uses to taste her. Next, he performs a species-specific wing song where he extends and vibrates one wing at a time towards her. Next, the male fly licks the genitalia of the female to open up the vaginal plate and further assess pheromonal cues, and lastly attempts copulation by curling up his abdomen (Ebbs and Amrein 2007). Mating and reproduction depends on the correct performance of this ritual. Each of these courtship behaviors is easily observable, and thus courtship makes for an ideal behavioral model.

We previously reported the identification of *Trapped in endoderm* 1 (*Tre1*) as a novel gene controlling courtship initiation in *Drosophila* (Luu *et al.* 2015). *Tre1* encodes an orphan Gprotein-coupled receptor (GPCR) that is required for both germ cell migration and establishment of cell polarity (Kunwar *et al.* 2008; Yoshiura *et al.* 2012), and is closely related to a family of GPCRs in vertebrates that respond to hormones and neurotransmitters, including melatonin and histamine (Yoshiura *et al.* 2012). We identified a set of neurons that express the transgene *Tre1-GAL4,* which, when genetically feminized either through expression of the female-specific splicing factor Transformer (Tra^F^), or expression of an RNAi construct targeted to the male-specific transcripts of the *fruitless* gene (*Fru^M^*), result in males displaying unusually rapid courtship initiation (Tran *et al.* 2014; Luu *et al.* 2015). We also found that feminization of *Tre1*-expressing cells led to a competitive reproductive advantage, and that *Tre1-GAL4* is expressed in a sexually dimorphic fashion in the olfactory organs and adult CNS (Luu *et al.* 2015).

Tre1 can signal through multiple heterotrimeric G-proteins. In establishment of cell polarity, the current model suggests that upon detection of an extrinsic signal, Tre1 activates the G-protein G_o_α, which then recruits Pins, which subsequently recruits the entire polarity complex (Yoshiura *et al.* 2012). In germ cell migration, Tre1 signals through the G-proteins Gγ1 and Gβ13f (Kunwar *et al.* 2008). Additionally, Tre1 directs the redistribution of the *D. melanogaster* E-cadherin, encoded by *shotgun (shg),* resulting in its polarized downregulation.

Here we further characterize the role of the Tre1 cells in courtship behavior. We confirm that *Tre1* function is required in neurons, and that genetic silencing of *Tre1-GAL4* expressing neurons results in rapid courtship initiation. We further identify histamine as the likely ligand of the Tre1 GPCR in courtship, and provide behavioral evidence for the negative regulation of *shotgun* by Tre1 during courtship. This is the first demonstration for a role of histamine in courtship behavior, as well as the first evidence for the existence of metabotropic histamine receptors in *Drosophila*.

## Materials and Methods

### Fly Strains and Genetics

Fly stocks were maintained at 25° C on standard cornmeal/ yeast/ molasses medium. With the exception of RNAi strains from the Transgenic RNAi Project (TRiP), all mutant alleles and transgenes were introgressed for five generations into our standard lab background [*w^1118^*; Wild Type Berlin (*w*; WTB)]. *UAS-Tra^F^*, *UAS-mCD8-GFP*, *UAS-shgRNAi*, *UAS-Gγ1RNAi*, UAS-*Gβ13fRNAi, UAS-ctaRNAi, Hdc^MB07212^*, *Hdc^JK910^*, *UAS-NaChBac*, and *UAS-Tre1RNAi* were all obtained from the Bloomington Drosophila Stock Center (Bloomington, Indiana) (stock nos. 4590, 5130, 32904, 25934, 31134, 31132, 25260, 64203, 9469, and 34956). *Tre1-GAL4* was isolated using meiotic recombination from the strain *GAL4^9-210^* (Luu *et al.* 2015). The strains *GAL4^9-210^*, *w^1118^*; *UAS-TNT(II)*, and *UAS-Elav-GAL4[3E1]* were gifts from Ulrike Heberlein.

### Courtship Assays

Courtship assays were performed according to published protocols (Villella *et al.* 1997). Virgin males were collected and kept in isolation for 1-4 days. After this period of isolation, each male was presented with a single 1-to-4-day-old *w^1118^*;WTB virgin female. Custom plexiglass chambers were used to contain the flies during the courtship assays, with each chamber being 10 mm in diameter and 6 mm in height. One male and one female were placed in each chamber, separated by a plastic transparency. After an acclimation period of 2-3 hours in the dark, plastic transparencies were removed to initiate contact between the pairs. Courtship behavior was recorded in infrared light for 20 minutes.

### Immunofluorescence

The CNS and peripheral tissue were dissected and fixed according to (Wu and Luo 2006) with the following modifications: tissues were fixed for 1 hr, incubated with NGS block for 24 hr, stained in primary antibody for 4 days, and secondary for 2 days. NC82 mouse anti-Brp was used at 1:50 (Developmental Studies Hybridoma Bank, AB 2314866). Rabbit anti-GFP was used at 1:750 (Life Technologies, A6455). Alexa Fluor 488 goat anti-rabbit was used at 1:1000 (Jackson Immunoresearch, 111-545-144). Alexa Fluor 594 goat anti-mouse was used at 1:1000 (Jackson Immunoresearch, 705-586-147). Images were taken using a Zeiss LSM700 confocal microscope.

### Statistics

An a level of 0.05 was used in all experiments. Data are represented as boxplots where the median is the middle line, and the 1st and 3rd quartiles are the lower and upper edges of the box, respectively. Whiskers represent the lowest and highest data points still within 1.5x interquartile range, while open dots represent data points outside of this range. Statistics were performed on log-transformed means of wing song latency. Data were analyzed using one-way analysis of variance (ANOVA), followed by Tukey’s HSD *post hoc* test. No statistical tests were used to predetermine sample sizes, but our sample sizes are consistent with those reported in previous publications (Tran *et al.* 2014; Luu *et al.* 2015).

### Reagent and Data Availability

All strains and reagents are available upon request. The full dataset will be uploaded as a supplementary file upon publication.

## Results

### *Tre1* is required in neurons for normal courtship behavior

As we have previously shown, (Luu *et al.* 2015), *Tre1* is required for normal courtship, and is expressed in neurons in both the periphery and CNS. To demonstrate that neuronal expression of *Tre1* is required for its role in courtship behavior, we used the pan-neuronal driver *elav-GAL4^c155^* to drive expression of *Tre1-RNAi* (Figure 1A). When we knocked down *Tre1* in neurons, male flies displayed rapid courtship initiation relative to background controls. Average time to courtship initiation in *elav-GAL4/Y; UAS-Tre1RNAi/+* males was 19 seconds, compared with 30-43 seconds in heterozygous genetic background controls (Figure 1A). These results demonstrate that *Tre1* expression is required specifically in neurons to establish a normal courtship initiation time.

**Figure 1.**
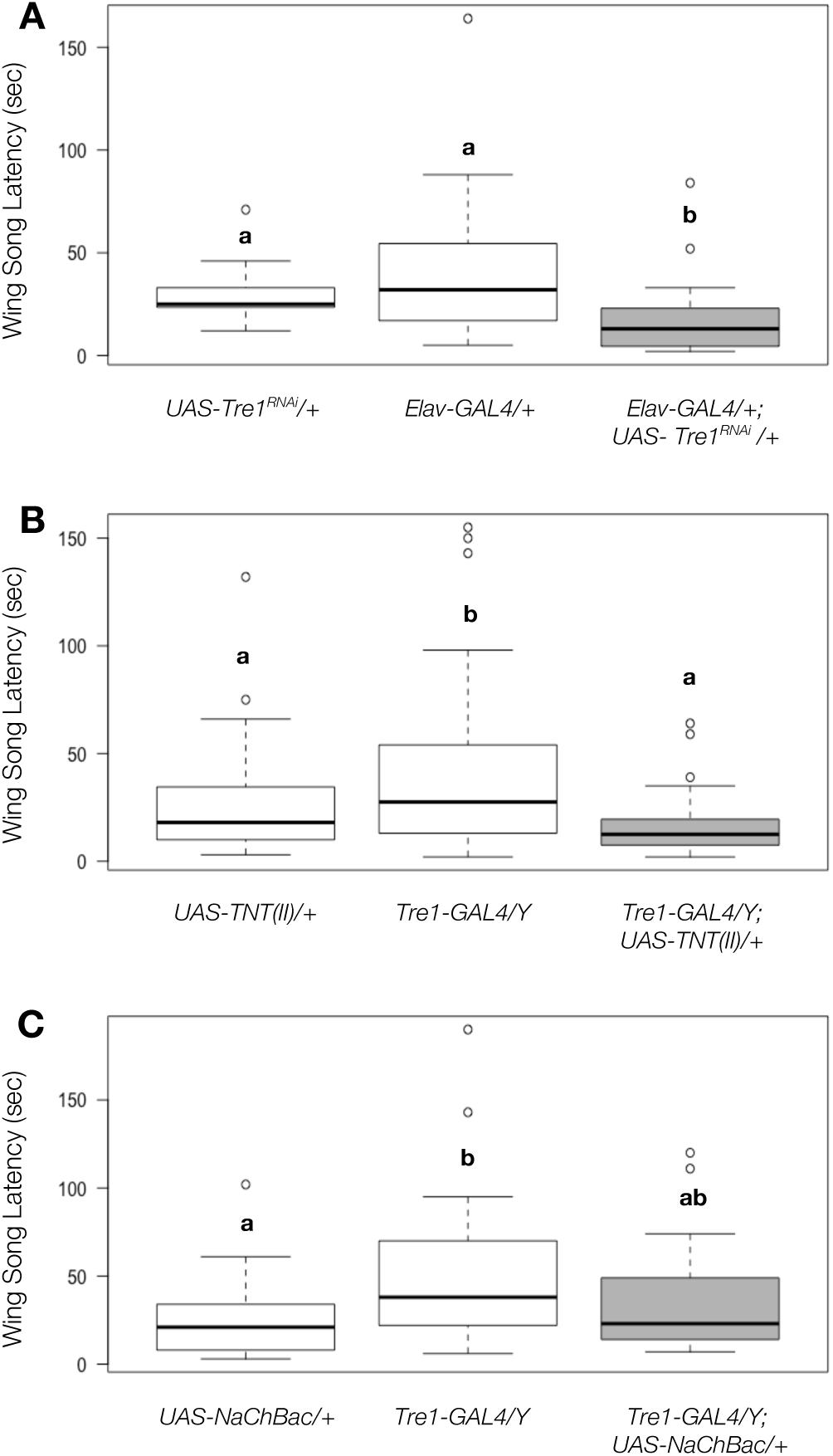
*Tre1* is required in neurons for normal courtship behavior and inactivation of Tre1 neurons results in rapid courtship initiation. **A.** *elav-GAL4/Y; UAS-Tre1RNAi*/+ males initiate courtship in 19 seconds, on average, compared with 30-43 seconds in controls (N=11-16, One-Way ANOVA with Tukey’s HSD posthoc analysis. **B.** *Tre1-GAL4/Y; UAS-TNT(II)/+* males initiate courtship in 17 seconds, on average, compared with 42 seconds in *Tre1-GAL4/+* controls and 25 seconds in *UAS-TNT/+* controls. (N=42-44, One-Way ANOVA with Tukey’s HSD posthoc analysis). **C.** Expression of *UAS-NaChBac* in *Tre1-GAL4* neurons has no effect on the speed of courtship initiation relative to controls (N=21-24, One-Way ANOVA with Tukey’s HSD posthoc analysis). Boxplots represent the median as the middle line, and the 1st and 3rd quartiles as the lower and upper edges, respectively. Whiskers represent the lowest and highest data points still within 1.5x interquartile range. Open dots represent outliers (data points outside 1.5x interquartile range). Boxes sharing the same letter do not differ significantly, while boxes with different letters are significantly different (p≤ 0.05).

### Silencing of *Tre1* cells results in rapid courtship initiation

We have previously shown that feminization of *Tre1* cells in males via expression of the female splicing factor Transformer results in rapid courtship initiation (Luu *et al.* 2015). This is an unusual phenotype, as feminization of neurons generally leads to delayed or absent courtship. We therefore considered the possibility that developmental feminization of the *Tre1* neurons results in an unusual gain-of-function effect that was not representative of the gene’s true function. To address this question, we used genetic techniques to examine the effects of silencing, and activation, of the *Tre1*-expressing neurons. Silencing the *Tre1-GAL4* neurons via expression of tetanus toxin (*UAS-TNT(II)*) resulted in males with wing song latency significantly shorter than that of *Tre1-GAL4*/Y background controls (average time to courtship initiation was 17 seconds for *Tre1-GAL4/Y; UAS-TNT(II)/+* and 42 seconds in *Tre1-GAL4/Y* controls) and these males were also somewhat faster than *UAS-TNT(II)/+* males (25 seconds to initiate courtship), though this result did not achieve statistical significance, due at least in part to the fact that *UAS-TNT(II)/+* males also display somewhat rapid courtship initiation (Fig. 1B). We attempted to silence the *Tre1-GAL4* neurons using a variety of alternative transgenes, but found that, in most cases, the result was lethality, or flies with profound locomotor defects that would impact courtship behavior in a nonspecific fashion (not shown).

In order to activate the *Tre1*-expressing neurons, we drove expression of a voltage-gated bacterial sodium channel, *UAS-NaChBac* (Hodge 2009) with *Tre1-GAL4*. There was no significant difference in wing song latency in *Tre1-GAL4/Y*; *UAS-NaChBac/+* males relative to either of the background controls (average time to courtship initiation for *Tre1-GAL4/Y*; *UAS-NaChBac/+* males was 36 seconds, compared with 27-52 seconds for heterozygous background controls; Figure 1C). These experiments suggest that feminizing the *Tre1* neurons results in loss of normal neuronal function, which is supported further by our previously-published experiments showing that a loss-of-function mutation in Tre1 (*Tre1^EP496^*) phenocopies feminization of the *Tre1* neurons (Luu *et al.* 2015).

### Feminization of *Tre1* cells results in the loss of *Tre1-GAL4* expression

We have previously shown that *Tre1-GAL4* is expressed in a sexually dimorphic pattern in the brain, with more expression in the male brain than the female brain (Luu *et al.* 2015). We hypothesized that feminization of the *Tre1*-expressing cells would lead to the loss of this sexually dimorphic expression pattern. To test this hypothesis, we used *Tre1*-GAL4 to drive simultaneous expression of UAS-*Tra^F^* and UAS-mCD8-*GFP* (Figure 2). Consistent with our hypothesis, *Tre1-GAL4/Y; UAS-Tra^F^/+; UAS-mCD8-GFP/+* males have little to no GFP expression compared with *Tre1-GAL4/Y; UAS-mCD8-GFP/+* males (Figures 2A, B). These results provide further evidence that feminization of the *Tre1*-expressing neurons results in a loss-of-function effect.

**Figure 2.**
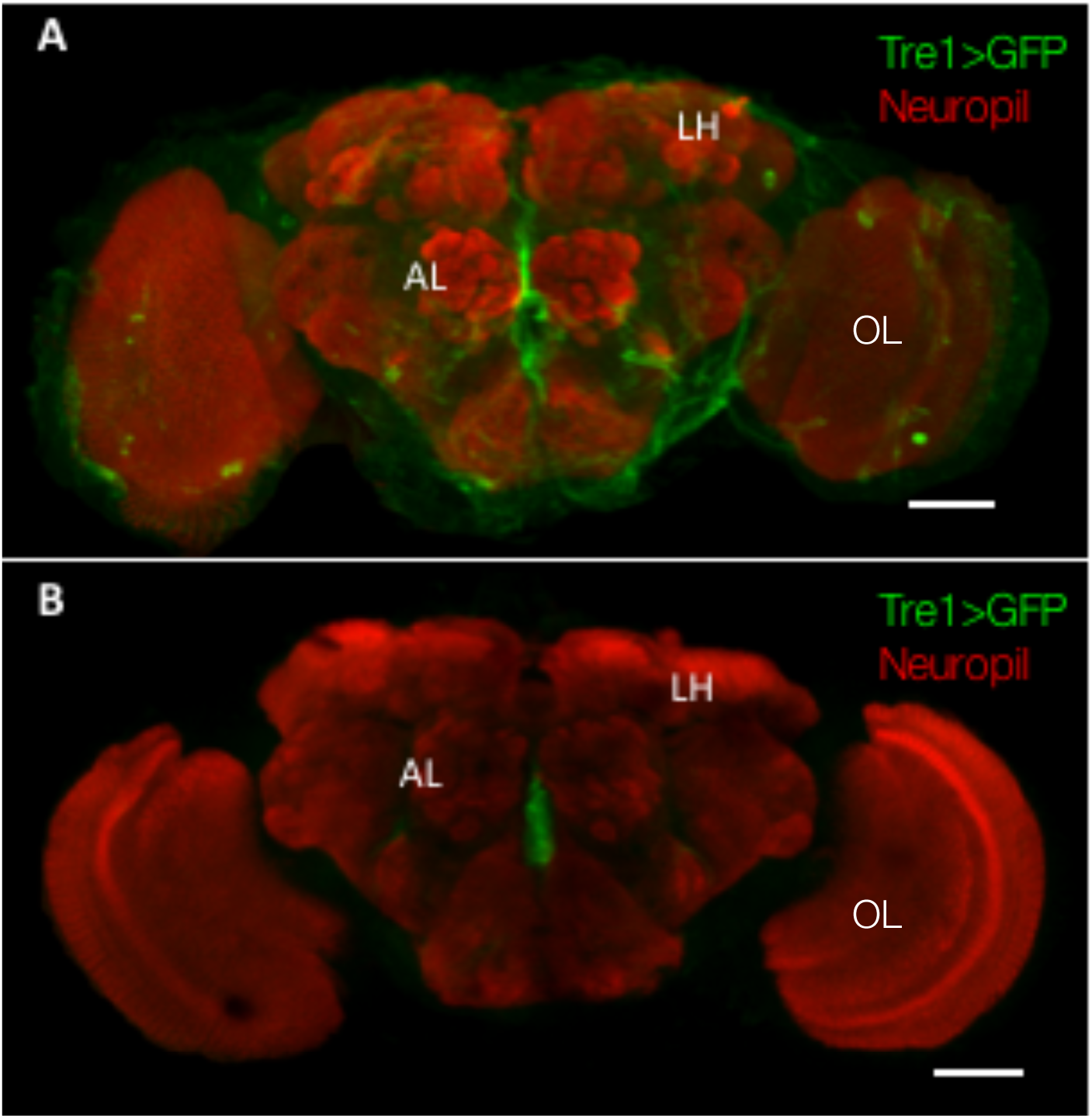
*Tre1-GAL4* expression in male brains is lost when *Tre1* cells are feminized. **A-B**. Representative confocal images of brains from male flies expressing GFP in *Tre1*-expressing cells; scale bars: 50 μM **A.** In *Tre1-GAL4>UAS-GFP* male brains, *Tre1* is expressed in a variety of regions, including the antennal lobe (AL), lateral horn (LH), and optic lobes (OL). **B.** In *Tre1-GAL4>UAS-Tra^F^*, *GFP* male brains, *Tre1-GAL4>UAS-GFP* expression is lost.

### Shotgun/E-cadherin is involved in regulating courtship initiation

The Tre1 GPCR has been studied in other contexts, and some downstream components of the G-protein signaling pathway are known in those systems (Kunwar *et al.* 2008; Yoshiura *et al.* 2012). To begin to characterize the signal transduction pathway downstream of Tre1 in courtship, we screened known downstream effectors of Tre1 for aberrant courtship initiation. Based on the findings of other groups, we tested a host of candidate genes including: *Gγ1*, *Gβ13f*, the *G*_α12,13_ subunit encoded by *concertina*, and the E-cadherin Shotgun; of these, only knockdown of *shotgun* resulted in aberrant courtship behavior (Supplementary Figure and Figure 3A).

**Figure 3.**
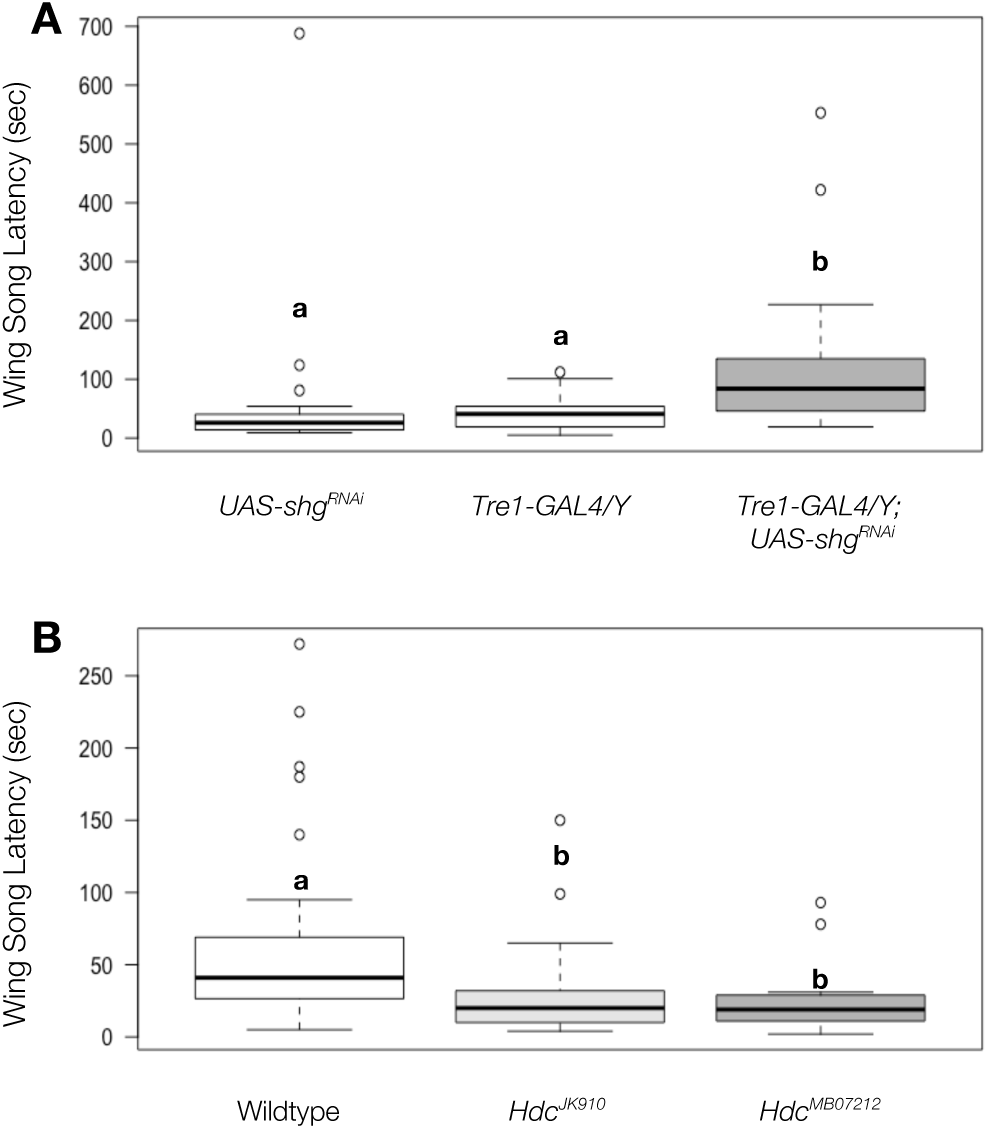
E-cadherin and histamine are involved in the timing of courtship initiation. **A.** *Tre1GAL4/Y;UAS-shg-RNAi/+* males display a significant delay in wing song latency (127 s) relative to control animals (43-59 s) (N=19-24, One-Way ANOVA with Tukey’s HSD post-hoc analysis). **B.** Histidine decarboxylase mutants, *Hdc^K910^ and Hdc^MB07212^*, display rapid wing song latency (31 s, n=25, P= 2.85 × 10^-3^ and 25 s, n= 21, P= 2.04 × 10^-3^, respectively) relative to control animals (61 s, n=39). Boxplots represent the median as the middle line, and the 1st and 3rd quartiles as the lower and upper edges, respectively. Whiskers represent the lowest and highest data points still within 1.5x interquartile range. Open dots represent outliers (data points outside 1.5x interquartile range). Boxes sharing the same letter do not differ significantly, while boxes with different letters are significantly different (p≤ 0.05).

*shotgun* (*shg*) encodes *D. melanogaster* E-cadherin, which is responsible for cell-cell adhesion, and mutations in *shg* result in the opposite effect of mutations in *Tre1* on germ cell migration (Kunwar *et al.* 2008). Knocking down *shg* in *Tre1* cells via expression of *shg-RNAi* resulted in a significantly slower time to courtship initiation compared with control males. *shg* knockdown males took, on average, 127 seconds to initiate courtship compared to 59 and 43 seconds in background controls (Fig. 3A). These results demonstrate that *shg*/*E-cad* is involved in the regulation of courtship initiation time, and suggest that E-cadherin is a downstream component regulated by *Tre1* signal transduction.

### Histamine is the likely ligand of the Tre1 receptor

Tre1 encodes an orphan G-protein-coupled receptor (GPCR), with an endogenous ligand that has yet to be identified. The Tre1 receptor shares sequence homology with a family of GPCRs that respond to hormones and neurotransmitters including melatonin and histamine, as well as being more distantly related to chemokine receptors such as CXCR4 (Kunwar *et al.* 2003) In order to identify the Tre1 ligand, we screened mutations in candidate genes for courtship phenotypes and found that male flies mutant for histidine decarboxylase (*Hdc*) displayed rapid courtship initiation (Figure 3B). Histidine decarboxylase catalyzes the decarboxylation of histidine to form the neurotransmitter histamine (Burg *et al.* 1993). We tested two alleles of *Hdc*, *Hdc^JK910^* and *Hdc^MB0721^*, and found that males homozygous for these alleles initiated courtship on average in 31 and 25 seconds, respectively, compared to 61 seconds in control flies. These data demonstrate that the neurotransmitter histamine is required for normal courtship behavior, and strongly suggest that histamine is the ligand for Tre1 in courtship initiation.

## Discussion

### *Tre1* is required in neurons for normal courtship behavior

We previously demonstrated that rapid courtship initiation results from both feminization of *Tre1-GAL4*-expressing cells as well as mutation of the GPCR-encoding *Tre1* gene itself (Luu *et al.* 2015). Here we demonstrate that this requirement is specific to neurons–when *Tre1* is knocked down specifically in neurons using the pan-neuronal driver *elav-GAL4^c155^*, males exhibit rapid courtship initiation (Figure 1A). In addition, we provide three lines of evidence that reducing the speed of courtship initiation is a normal function of the *Tre1*-expressing neurons: first, loss-of-function mutations in *Tre1* lead to rapid courtship (Luu *et al.* 2015), suggesting that rapid courtship initiation is a result of loss of normal neuronal function. Second, silencing the *Tre1-GAL4* neurons through expression of tetanus toxin results in rapid courtship initiation, similar to that seen with mutation of *Tre1* or feminization of the *Tre1* neurons (Figure 1B). Third, we show that feminizing the *Tre1-GAL4* neurons results in loss of Tre1 expression (Figure 2B), further suggesting that feminization of the *Tre1-GAL4* neurons results in a reduction of normal function.

While silencing of the *Tre1-GAL4* neurons resulted in significantly rapid courtship initiation relative to the *Tre1-GAL4/Y* genetic control, the decrease in courtship initiation speed when compared to the *UAS-TNT(II)/+* control did not achieve statistical significance (Figure 1B). We have three hypotheses to explain this result. First, it is possible that this is due to the fact that we were forced to use a relatively “weak” combination of transgenes for this experiment. We tried a number of approaches to silencing or eliminating the *Tre1-GAL4* neurons, including alternative insertions of the *UAS-TNT* transgene and expression of the cell death protein Reaper (Rpr). We found that, with the exception of the second-chromosomal insertion of *UAS-TNT*, these manipulations resulted in lethality, or, when flies survived, profound locomotor deficiencies. Thus, the experiment may have failed to achieve significance due simply to the relatively weak silencing that was possible with the available tools.

Additionally, the time to courtship initiation for the *UAS-TNT(II)/+* controls was relatively rapid compared with typical control speeds, which may contribute to “masking” the effect of silencing the *Tre1-GAL4* cells relative to that control.

Finally, it is a formal possibility that, while *Tre1* gene function is required for the timing of courtship initiation, the *activity* of the *Tre1*-expressing cells is not required for the behavior itself. In such a scenario, there might be a requirement for *Tre1* in directing the development of a neural circuit, such that the *Tre1* neurons influence the development of another group of cells, but are not, themselves, required in the adult brain for the expression of the behavior. We are currently working to test this hypothesis, first, by establishing whether there is a developmental requirement for *Tre1*.

### Histamine Is Required for Courtship Behavior

In order to investigate possible ligands for the Tre1 receptor, we screened mutations in candidate genes for courtship phenotypes. Because Tre1 has significant sequence similarity to vertebrate histamine receptors (Kunwar *et al.* 2003), we tested flies mutant for *histidine decarboxylase* (*Hdc*). Male flies homozygous for loss-of-function mutations in *Hdc* display the same unusually rapid courtship initiation phenotype we observe in *Tre1* hypomorphs and *Tre1-GAL4/Y;UAS-Tra^F^*/+ males (Figure 3B). Further, histamine is the primary neurotransmitter responsible for photoreception in insects (Elias and Evans 1983; Hardie 1987; Melzig *et al.* 1996), and Tre1-GAL4 is expressed in the optic lobe in both male and female flies (Figure 2A and data not shown). These data strongly suggest that histamine is the Tre1 ligand.

In addition to the requirement for histamine in insect photoreception, histamine is involved in mechanosensation and thermotaxis in flies (Buchner *et al.* 1993; Melzig *et al.* 1996; Hong *et al.* 2006), as well as regulation of sleep-wake cycles in both flies and mammals (Wauquier *et al.* 1981; Thakkar 2011; Oh *et al.* 2013).

There are four known metabotropic histamine receptors (H1-H4) in humans, all of which are G-protein-coupled receptors, and these regulate various biological processes including: sleep-wake regulation, immune responses, neurotransmission, and chemotaxis (Brown *et al.* 2001; Panula *et al.* 2015). This work is the first demonstration of histamine’s involvement in courtship behavior; moreover, the only characterized receptors for histamine in flies are histamine-gated chloride channels (Zheng *et al.* 2002)—ours is the first suggestion of a metabotropic histamine receptor in *Drosophila*, and only the second in invertebrates (Roeder 2003).

### The Role of E-cadherin In Courtship Behavior

Males in which E-cadherin has been knocked down in the *Tre1-GAL4* neurons display a significant delay in courtship initiation, the opposite phenotype from that seen with loss of *Tre1* Figure 3A). To our knowledge, this is the first demonstration for a role of E-cadherin in courtship regulation.

In germ cell migration, *Tre1* expression results in polarized downregulation of Shg/E-cad, and this downregulation is necessary for germ cell dispersal (but not sufficient for transepithelial migration) (Kunwar *et al.* 2008). Thus, when germ cells were mutant for *Tre1*, they failed to disperse due to continued expression of E-cad, while, when E-cad was mutated, germ cell dispersal occurred prematurely. Moreover, downregulation of E-cadherin promotes neuronal migration in mice (Itoh *et al.* 2013; Chen *et al.* 2015).

E-cadherin is expressed widely in both vertebrate and *Drosophila* nervous systems (Fannon and Colman 1996; Uchida *et al.* 1996; Iwai *et al.* 1997; BrusÉS 2000; Tepass *et al.* 2000; Dumstrei *et al.* 2003; Prakash *et al.* 2005; Fung *et al.* 2008), and cadherins (both N- and E-cadherin) have been implicated in axon targeting (Prakash *et al.* 2005), synaptic partner recognition (Shapiro and Colman 1999), synaptogenesis (Benson and Tanaka 1998; Takai *et al.* 2003; Bozdagi *et al.* 2004; Arikkath and Reichardt 2008), synaptic adhesion (Fannon and Colman 1996; BrusÉS 2000), and synaptic plasticity (Tang *et al.* 1998; Arikkath and Reichardt 2008).

Given our data that loss of both histamine and Tre1 lead to rapid courtship initiation, while *shg/E-cad* mutants have the opposite phenotype, we propose that histamine signals through Tre1 to downregulate E-cad, and that this interaction is required to properly regulate either synaptogenesis or synaptic adhesion between the Tre1 neurons and their targets

Both E- and N-cadherin-containing synapses are present throughout the CNS of the adult mouse, and many synapses label with neither antiserum, suggesting that still other cadherins are present at those junctions. Further, the “zones” of E- and N-cadherin expression are largely nonoverlapping, and it has been proposed that cadherins serve to organize the specificity of synaptic junctions as well as provide adhesive connections between neurons (Fannon and Colman 1996). N-cadherin is found at both the presynaptic and postsynaptic terminals in cultured rat hippocampal neurons, and clusters together with synaptic markers at developing synapses (Benson and Tanaka 1998). Later, however, N-cadherin is retained at excitatory synapses, but is lost from inhibitory synapses, while E-cadherin is often found at inhibitory synapses (Benson and Tanaka 1998; BrusÉS 2000), suggesting that synapse formation and maintenance may be dependent on developmental regulation and expression of specific types of cadherins.

We propose a model in which the Tre1 neurons modulate the speed of courtship initiation, in response to an as-yet-unidentified external cue. Histamine, signaling through the Tre1 receptor, reduces the expression of E-cadherin, and this is necessary for establishment of the correct number (or strength) of synaptic connections between the Tre1 neurons and downstream neurons in the courtship circuit. In neurons deficient for either histamine or Tre1, inappropriate expression of E-cad causes increased numbers or strength of E-cad-containing synaptic connections, and this imbalance results in rapid courtship initiation. Conversely, when *shg* is mutant, the number or strength of E-cad-containing synapses between Tre1 neurons and their downstream partners is decreased, leading to delayed or absent courtship initiation

## Conclusions

We have shown that the GPCR Tre1 is required in neurons to regulate the speed of courtship initiation, and that silencing of Tre1-expressing neurons may lead to rapid courtship initiation. We have also established a role for both E-cadherin and histamine in courtship behavior, and, importantly, have provided the first evidence for metabotropic histamine receptors in *Drosophila*, and, indeed, only the second example of metabotropic histamine receptors in invertebrates. Ongoing and future experiments will further investigate the role of histamine signaling in courtship behavior, as well as examine the expression of E-cad in the *Tre1* neurons and the effects of mutation of *Tre1* and *Hdc* on E-cad expression and synapse formation and distribution.

## Literature Cited

Arikkath J., Reichardt L. F., 2008 Cadherins and catenins at synapses: roles in synaptogenesis and synaptic plasticity. Trends Neurosci. 31: 487–94.

Benson D. L., Tanaka H., 1998 N-cadherin redistribution during synaptogenesis in hippocampal neurons. J. Neurosci. 18: 6892–904.

Bozdagi O., Valcin M., Poskanzer K., Tanaka H., Benson D. L., 2004 Temporally distinct demands for classic cadherins in synapse formation and maturation. Mol. Cell. Neurosci. 27: 509–21.

Brown R. E., Stevens D. R., Haas H. L., 2001 The physiology of brain histamine. Prog. Neurobiol. 63: 637–72.

BrusÉS J. L., 2000 Cadherin-mediated adhesion at the interneuronal synapse. Curr. Opin. Cell Biol. 12: 593–7.

Buchner E., Buchner S., Burg M. G., Hofbauer A., Pak W. L., Pollack I., 1993 Histamine is a major mechanosensory neurotransmitter candidate in Drosophila melanogaster. Cell Tissue Res. 273: 119–25.

Burg M. G., Sarthy P. V., Koliantz G., Pak W. L., 1993 Genetic and molecular identification of a Drosophila histidine decarboxylase gene required in photoreceptor transmitter synthesis. EMBO J. 12: 911–9.

Chen D., Wu Z., Luo L.-J. J., Huang X., Qian W.-Q. Q., Wang H., Li S.-H. H., Liu J., 2015 E-cadherin maintains the activity of neural stem cells and inhibits the migration. Int J Clin Exp Pathol 8: 14247–51.

Dumstrei K., Wang F., Hartenstein V., 2003 Role of DE-cadherin in neuroblast proliferation, neural morphogenesis, and axon tract formation in Drosophila larval brain development. J. Neurosci. 23: 3325–35.

Ebbs M. L., Amrein H., 2007 Taste and pheromone perception in the fruit fly Drosophila melanogaster. Pflugers Arch. 454: 735–47.

Elias M. S., Evans P. D., 1983 Histamine in the insect nervous system: distribution, synthesis and metabolism. J. Neurochem. 41: 562–8.

Fannon A. M., Colman D. R., 1996 A model for central synaptic junctional complex formation based on the differential adhesive specificities of the cadherins. Neuron 17: 423–34.

Fung S., Wang F., Chase M., Godt D., Hartenstein V., 2008 Expression profile of the cadherin family in the developing Drosophila brain. J. Comp. Neurol. 506: 469–88.

Hardie R. C., 1987 Is histamine a neurotransmitter in insect photoreceptors? J. Comp. Physiol. A 161: 201–13.

Hodge J. J., 2009 Ion channels to inactivate neurons in Drosophila. Front Mol Neurosci 2: 13.

Hong S.-T. T., Bang S., Paik D., Kang J., Hwang S., Jeon K., Chun B., Hyun S., Lee Y., Kim J., 2006 Histamine and its receptors modulate temperature-preference behaviors in Drosophila. J. Neurosci. 26: 7245–56.

Itoh Y., Moriyama Y., Hasegawa T., Endo T. A., Toyoda T., Gotoh Y., 2013 Scratch regulates neuronal migration onset via an epithelial-mesenchymal transition-like mechanism. Nat. Neurosci. 16: 416–25.

Iwai Y., Usui T., Hirano S., Steward R., Takeichi M., Uemura T., 1997 Axon patterning requires DN-cadherin, a novel neuronal adhesion receptor, in the Drosophila embryonic CNS. Neuron 19: 77–89.

Kunwar P., Sano H., Renault A., Barbosa V., Fuse N., Lehmann R., 2008 Tre1 GPCR initiates germ cell transepithelial migration by regulating Drosophila melanogaster E-cadherin. J Cell Biology 183:157–168.

Kunwar P. S., Starz-Gaiano M., Bainton R. J., Heberlein U., Lehmann R., 2003 Tre1, a G protein-coupled receptor, directs transepithelial migration of Drosophila germ cells. PLoS Biol. 1: E80.

Luu P., Zaki S., Tran D., French R., 2015 A Novel Gene Controlling the Timing of Courtship Initiation in Drosophila melanogaster. Genetics 202: 1043–1053.

Melzig J., Buchner S., Wiebel F., Wolf R., Burg M., Pak W. L., Buchner E., 1996 Genetic depletion of histamine from the nervous system of Drosophila eliminates specific visual and mechanosensory behavior. J. Comp. Physiol. A 179: 763–73.

Oh Y., Jang D., Sonn J. Y., Choe J., 2013 Histamine-HisCl1 receptor axis regulates wake-promoting signals in Drosophila melanogaster. PLoS ONE 8: e68269.

Panula P., Chazot P. L., Cowart M., Gutzmer R., Leurs R., Liu W. L., Stark H., Thurmond R. L., Haas H. L., 2015 International Union of Basic and Clinical Pharmacology. XCVIII. Histamine Receptors. Pharmacol. Rev. 67: 601–55.

Prakash S., Caldwell J. C., Eberl D. F., Clandinin T. R., 2005 Drosophila N-cadherin mediates an attractive interaction between photoreceptor axons and their targets. Nat. Neurosci. 8: 443–50.

Roeder T., 2003 Metabotropic histamine receptors—nothing for invertebrates? Eur. J. Pharmacol. 466: 85–90.

Shapiro L., Colman D. R., 1999 The diversity of cadherins and implications for a synaptic adhesive code in the CNS. Neuron 23: 427–30.

Takai Y., Shimizu K., Ohtsuka T., 2003 The roles of cadherins and nectins in interneuronal synapse formation. Curr Opin Neurobiol 13: 520–526.

Tang L., Hung C. P., Schuman E. M., 1998 A role for the cadherin family of cell adhesion molecules in hippocampal long-term potentiation. Neuron 20: 1165–75.

Tepass U., Truong K., Godt D., Ikura M., Peifer M., 2000 Cadherins in embryonic and neural morphogenesis. Nat. Rev. Mol. Cell Biol. 1: 91–100.

Thakkar M. M., 2011 Histamine in the regulation of wakefulness. Sleep Med Rev 15: 65–74.

Tran D. H., Meissner G. W., French R. L., Baker B. S., 2014 A small subset of fruitless subesophageal neurons modulate early courtship in Drosophila. PLoS ONE 9: e95472.

Uchida N., Honjo Y., Johnson K. R., Wheelock M. J., Takeichi M., 1996 The catenin/cadherin adhesion system is localized in synaptic junctions bordering transmitter release zones. J. Cell Biol. 135: 767–79.

Villella A., Gailey D. A., Berwald B., Ohshima S., Barnes P. T., Hall J. C., 1997 Extended reproductive roles of the fruitless gene in Drosophila melanogaster revealed by behavioral analysis of new fru mutants. Genetics 147: 1107–30.

Wauquier A., Broeck W. A. VAN DEN, Awouters F., Janssen P. A., 1981 A comparison between astemizole and other antihistamines on sleep-wakefulness cycles in dogs. Neuropharmacology 20: 853–9.

Wu J. S., Luo L., 2006 A protocol for dissecting Drosophila melanogaster brains for live imaging or immunostaining. Nat Protoc 1: 2110–5.

Yoshiura S., Ohta N., Matsuzaki F., 2012 Tre1 GPCR signaling orients stem cell divisions in the Drosophila central nervous system. Dev. Cell 22: 79–91.

Zheng Y., Hirschberg B., Yuan J., Wang A. P., Hunt D. C., Ludmerer S. W., Schmatz D. M., Cully D. F., 2002 Identification of two novel Drosophila melanogaster histamine-gated chloride channel subunits expressed in the eye. J. Biol. Chem. 277: 2000–5.

